# Cellular electrical impedance to profile SARS-CoV-2 fusion inhibitors and to assess the fusogenic potential of spike mutants

**DOI:** 10.1101/2022.12.13.520307

**Authors:** Emiel Vanhulle, Jordi Doijen, Joren Stroobants, Becky Provinciael, Sam Noppen, Dominique Schols, Annelies Stevaert, Kurt Vermeire

## Abstract

Despite the vaccination campaigns for COVID-19, we still cannot control the spread of SARS-CoV-2, as evidenced by the ongoing circulation of the Omicron variants of concern. This highlights the need for broad-spectrum antivirals to further combat COVID-19 and to be prepared for a new pandemic with a (re-)emerging coronavirus. An interesting target for antiviral drug development is the fusion of the viral envelope with host cell membranes, a crucial early step in the replication cycle of enveloped viruses. In this study, we explored the use of cellular electrical impedance (CEI) to quantitatively monitor morphological changes in real time, resulting from cell-cell fusion elicited by SARS-CoV-2 spike. The impedance signal in CEI-quantified cell-cell fusion correlated with the expression level of SARS-CoV-2 spike in transfected HEK293T cells. For antiviral assessment, we validated the CEI assay with the fusion inhibitor EK1 and measured a concentration-dependent inhibition of SARS-CoV-2 spike mediated cell-cell fusion (IC_50_ value of 0.13 μM). In addition, CEI was used to confirm the fusion inhibitory activity of the carbohydrate-binding plant lectin UDA against SARS-CoV-2 (IC_50_ value of 0.55 μM), which complements prior in-house profiling activities. Finally, we explored the utility of CEI in quantifying the fusogenic potential of mutant spike proteins and in comparing the fusion efficiency of SARS-CoV-2 variants of concern. In summary, we demonstrate that CEI is a powerful and sensitive technology that can be applied to studying the fusion process of SARS-CoV-2 and to screening and characterizing fusion inhibitors in a label-free and non-invasive manner.

**Importance:** Despite the success of the vaccines against SARS-CoV-2, new variants of the virus are still emerging and spreading, underlining the need for additional effective antiviral countermeasures. An interesting antiviral target for enveloped viruses is the fusion of the viral envelope with host cell membranes, a crucial early step in the life cycle of coronaviruses like SARS-CoV-2. Here, we present a sensitive impedance-based method to monitor in real-time cell-cell fusion elicited by the SARS-CoV-2 spike protein. With this technique we can profile entry inhibitors and determine the inhibitory potential of fusion inhibitors for SARS-CoV-2. In addition, with cellular electrical impedance we can evaluate the fusogenic properties of new emerging SARS-CoV-2 variants. Overall, the impedance technology adds valuable information on the fusion process of circulating coronaviruses and helps unravel the mode of action of new antivirals, opening new avenues for the development of next generation fusion inhibitors with improved antiviral activity.

## Introduction

Despite the successful COVID-19 vaccination campaign, we are still unable to control the spread of new variants and/or prevent re-infections. This underlines the need to continue with the development of effective antiviral compounds against SARS-CoV-2. An appealing target for drug development is the fusion of SARS-CoV-2 viral envelope with host cell membranes, an essential early step in the coronavirus life cycle.

Entry of SARS-CoV-2 in target cells involves several sequential steps that are mediated by the spike (S) protein that drives the fusion process by a series of coordinated conformational changes (1-4). The S protein is cleaved into two subunits: the S1 subunit, which recognizes the angiotensin-converting enzyme 2 (ACE2) through binding via its receptor-binding domain (RBD) (5, 6), and the S2 subunit, which harbors the fusion machinery. Generally, these two subunits are post-translationally cleaved at the S1-S2 site of the S protein by the host serine protease furin (7, 8), however, it is still debatable if the Omicron S protein is less cleavable by furin (9-11). After binding to human ACE2, a transmembrane receptor that is highly expressed in lung epithelial cells, a second cleavage event at the S2′ site must take place to render the S protein fully fusion-competent. For the initial SARS-CoV-2 variants (e.g., the Wuhan-Hu-1 strain), S2′ cleavage occurs preferably at the cell surface by type II transmembrane serine proteases (TTSPs) such as transmembrane protease serine 2 (TMPRSS2) (12, 13). Alternative to the furin/TMPRSS2 proteolytic activation, SARS-CoV-2 can enter via endocytosis with cathepsin B or L (CTSB/L) cleaving the S protein (9), an entry route proposed for the recent Omicron variants (9, 10). Either way, receptor binding and S2′ cleavage result in the formation of an elongated intermediate spike protein with its hydrophobic fusion peptide (FP) exposed, followed by the insertion of the activated fusion protein into the target host membrane (**Figure 1A**) (14). The subsequent collapse of the metastable spike intermediate into an energetically stable (low energy) hairpin-like configuration, the so-called 6-helix bundle complex of the S2 trimer, brings the viral and cellular membrane in close proximity for their merger and the completion of membrane fusion. Finally, the viral genome is released into the cytosol of the host cell to initiate the replication cycle.

**FIG 1.**
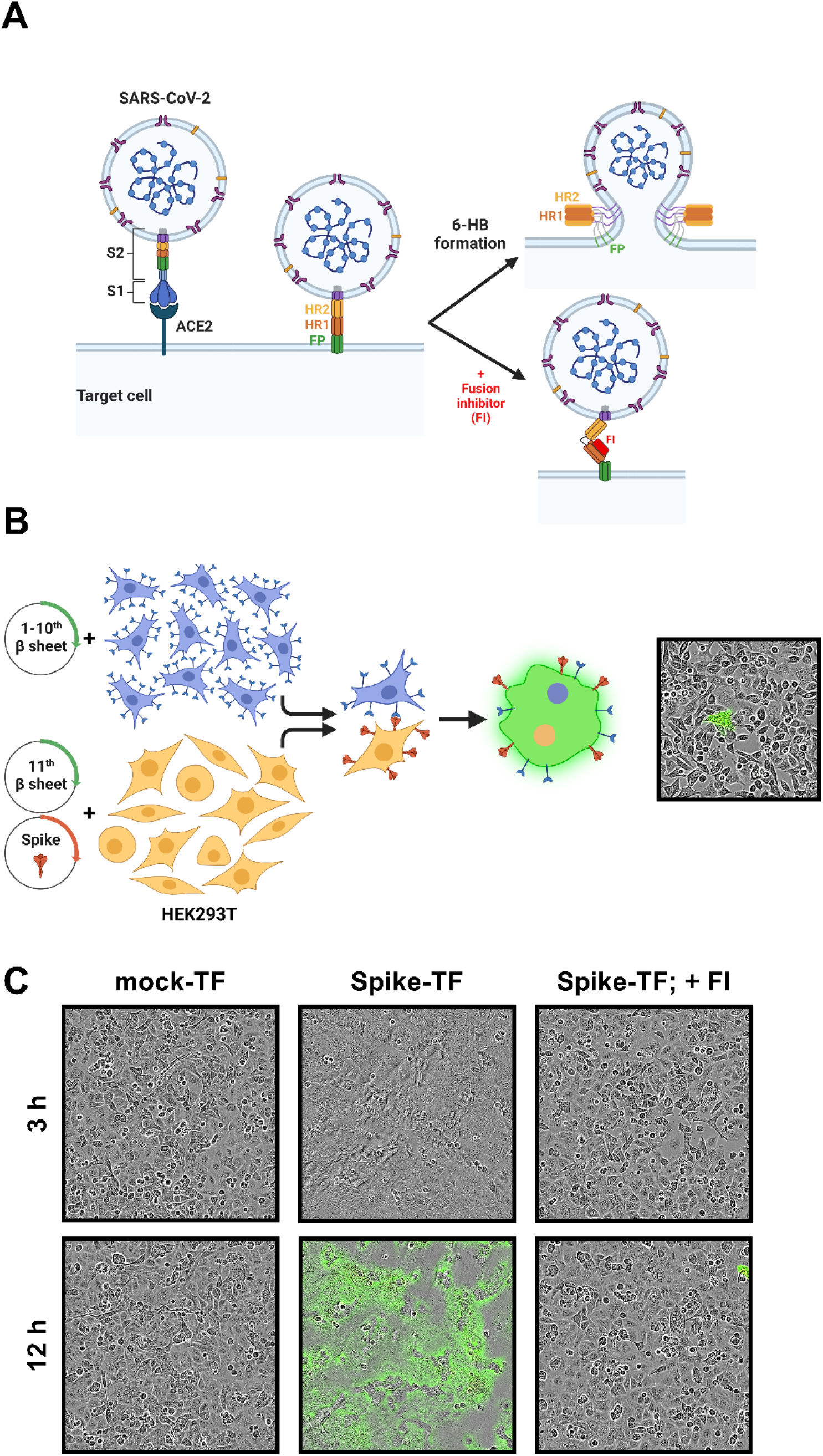
SARS-CoV-2 spike-transfected cells mimic viral envelope for fusion with ACE2^+^ target cell membrane. (**A**) Schematic representation of the fusion process for SARS-CoV-2 and a potential target for fusion inhibitors. See text for detailed description of the fusion process. FP, fusion peptide; HR, heptad repeat domain; 6-HB, 6 helix bundle; FI, fusion inhibitor. (**B**) Schematic representation of a split neongreen fusion assay (figure adapted from (23)). A549.ACE2^+^ cells (transfected to express the first 10 betasheets of neongreen) were overlayed with HEK293T cells co-transfected with a plasmid encoding the SARS-CoV-2 spike protein and a plasmid encoding the 11^th^ betasheet of neongreen. Only cell-cell fusion of an A549 cell with a HEK293T cell will result in the assembly of a functional neongreen protein and give a green fluorescence signal as the former expresses spike and the latter human ACE2. Light microscopic picture shows fused cells with neongeen expression (20x magnification). (**C**) Same as in (B). A549.ACE2^+^ cells were overlayed with HEK293T cells either transfected (TF) with an empty vector (left panels; mock-TF), or with Wuhan-Hu-1 S protein and left untreated (middle) or treated with the fusion inhibitor EK1 (2 μM; right panels). Light microscopic pictures were taken at 3 and 12 hours post overlay (20x magnification). Note that cell-cell fusion in the untreated spike-transfected condition is already visible at 3h post overlay but that neongreen fluorescence is still absent. Cartoons were created with BioRender (www.biorender.com).

Cellular electrical impedance (CEI) is a label-free, quantitative analytical method used to study cell morphological changes in real time. When an electrical circuit is applied to a cell monolayer grown in microtiter plates embedded with gold electrodes (15, 16), the continuous sweeping of non-invasive alternating current (AC) voltages over a predefined set of frequencies allows us to measure the impedance (Z) on this current. Cell Index (CI) is a quantitative measure of the status of the cells in an electrode-containing well and is based on the measured cell-electrode impedance (17). When cells are attached to and spread out over the electrodes, they act as insulating particles that will resist or impede the flow of the current. As a result, the impedance of the system (and its CI value) will increase when the cell monolayer becomes more confluent. On the other hand, when cells of a confluent cell culture die and lyse, the disruption of the cell monolayer facilitates current flow, which translates in a decrease in impedance over time. CEI has gained popularity in recent years as a method to monitor dynamic responses of cells. Among other areas of research, CEI has proven effective for cytotoxicity measurements and signalling pathway studies (18, 19), and is recently gaining popularity for virological studies as well (20, 21). For instance, CEI has been proposed as a generic screening method for fusion inhibitors targeting respiratory syncytial virus (RSV), dengue virus (DENV) and vesicular stomatitis virus (VSV), which are representatives of three viral fusion classes within the enveloped viruses (22).

Since the fusion of viral and cellular membranes induces changes in cell morphology, we explored the feasibility of CEI to monitor the fusion process of SARS-CoV-2 in real time in a cell-to-cell fusion format. In this work we (i) successfully optimized and validated an impedance-based fusion assay to quantifiable measure SARS-CoV-2 spike-induced cell-cell fusion, (ii) evaluated potential SARS-CoV-2 fusion inhibitors, and, (iii) compared the fusogenic potential of SARS-CoV-2 variants of concern (VOCs).

## Results

### SARS-CoV-2 spike-induced cell-cell fusion

In previous studies (23, 24), we demonstrated that SARS-CoV-2 viral entry can be mimicked by a cell-cell fusion system in which acceptor cells express the cellular human ACE2 receptor and donor cells the complementary viral spike protein. In our current work, the lung epithelial cell line A549 (stably transduced with human ACE2 to elevate the endogenous receptor level) was chosen as acceptor cell, whereas spike-transfected HEK293T cells were selected as donor cells because of the high plasmid transfection efficiency of the latter. Co-cultivation of both cell types results in fusion of the cells with the generation of multinucleated giant cells, the so-called syncytia that can be microscopically observed.

To better visualize this cell-cell fusion event, we previously designed and reported a split neon green assay (**Figure 1B**) (23). Real time microscopy can be used to monitor the fusion event (see also **Supplementary movie 1**) and to evaluate the inhibitory effect of a fusion inhibitor (**Figure 1C**). However, as the read-out of this assay is the fluorescence of the neongreen protein, a certain delay has to be taken into account between the initial cell-cell membrane fusion and the ultimate formation of an active fluorescent protein, which largely depends on the intermingling of the cytosolic content of both cells and the migration and assembly of both neongreen subunits (**Figure 1C**; compare 3h with 12h condition for untreated spike-transfected cells). Furthermore, when a single spike-transfected cell fuses with multiple ACE2-expressing cells (or vice versa), the fluorescent signal might not accurately reflect the number of fusion events because of a possible imbalance in the amount of the two neongreen subunits in the cytosol of the multinucleated cell. Therefore, an alternative fusion assay based on cellular electrical impedance (CEI) was explored.

### Cellular electrical impedance as a measure for SARS-CoV-2 spike-induced cell-cell fusion

In order to measure CEI, experiments are performed in specialized impedance E-plates, i.e., microtiter plates with embedded golden microelectrodes. A549.ACE2^+^ cells are first seeded into E-plates to obtain a confluent cell monolayer. When monitoring the impedance signal of this growing monolayer, a slow but steady increase in the cell index (CI) was observed during the initial adherence phase (**Figure 2**, grey line). At 24h post plating, the A549.ACE2^+^ cells are overlayed with HEK293T cells either transfected with empty vector (mock) or with a vector encoding the SARS-CoV-2 spike protein (**Figure 2**; phase #1). Cell-cell fusion is then triggered by the interaction of the viral fusion glycoproteins at the HEK293T cell surface with ACE2 receptor molecules on a neighbouring A549.ACE2^+^ cell. The CI value of the A549.ACE2^+^ cells exposed to mock-transfected control HEK293T cells slowly increases (because of residual cell proliferation and/or maturation of the cell layer, *i*.*e*., more compact cell-to-cell contacts) until a nearly steady state is reached (**Figure 2**, grey line). However, when spike-transfected HEK293T cells are added on top of the A549.ACE2^+^ cells, fusion already manifests within a few hours of cell overlay and the CI value sharply increases as a result of the generated syncytia (**Figure 2**, blue line). When cells fuse, the electrical current can no longer pass in between them because of a decrease in cell-cell borders and tight junctions in the cell monolayer. This CI increase correlates nicely with the formation of syncytia as determined by microscopy (see **Supplementary movie 2**). Once a maximum in cell-cell fusion is reached (**Figure 2**, blue line; phase #2), the CI value starts to decline because of the instability of the multinucleated cell membrane, and the subsequent cell lysis and destruction of the cell monolayer. At 24h post overlay, the cell monolayer is completely destroyed and the CI has returned to baseline level (**Figure 2**, blue line; phase #3).

**FIG 2.**
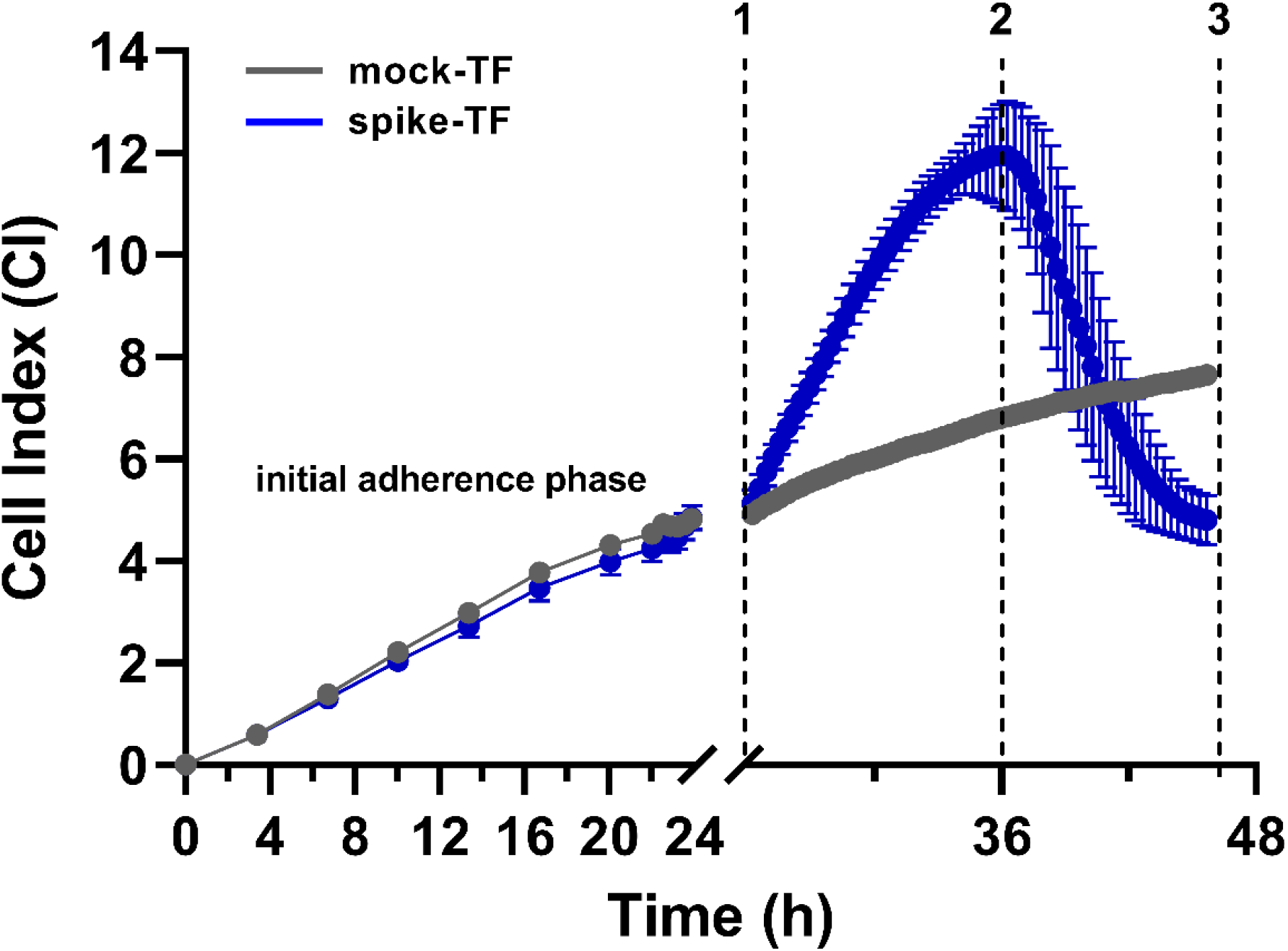
Comparison of impedance signal of A549.ACE2^+^ cells overlayed with mock-transfected versus Wuhan-Hu-1 SARS-CoV-2 spike-transfected HEK293T cells. At time point 0, A549.ACE2^+^ cells were seeded and impedance was recorded of the proliferating cell monolayer. At 24h post plating (phase #1), empty vector- (grey) and spike-transfected (blue) HEK293T cells were added. The graph depicts the raw impedance signal (expressed as cell index) over time of 4 technical replicates (mean ± SD). Vertical dotted lines 1 to 3 indicate important phases, which are further explained in the text. Note the bigger variation in CI response between the replicates during the disruption of the cell monolayer (starting at phase #2).

### Optimization of CEI for SARS-CoV-2 spike induced cell-cell fusion

In a first set of optimization experiments, different parameters of our CEI assay were investigated to reach optimal fusion and to obtain higher CI values. As shown in **Figure 3A**, the kinetics of cell-cell fusion depended on the ratio of acceptor:donor cells. Generally, an excess of spike-transfected donor cells enhanced the fusion process. An acceptor:donor cell ratio of 1:1 resulted in a nice response and was selected for further experiments (**Figure 3A**, blue curve). As expected, cell-cell fusion and the related impedance response relied on the complementary expression of ACE2 and S. This was evidenced by the absence of fusion either when A549.ACE2^+^ cells were combined with mock-transfected HEK293T cells, or when native A549 control cells (with low endogenous ACE2 levels) were combined with S-expressing HEK293T cells (**Supplementary Figure 1A**). Furthermore, trypsinization of the transfected HEK293T cells (as compared to collection of the cells by simply resuspending the easy detaching cell monolayer) had a positive outcome on cell-cell fusion, with a faster and more uniformily response (**Supplementary Figure 1B**). Lowering the incubation temperature of the HEK293T cells from 37 to 34°C during the spike biogenesis after transfection did not impact the initial phase of fusion but appears to somewhat reduce the maximum response (**Supplementary Figure 1C**). Importantly, the expression level of spike on the HEK293T cells was a crucial determinant for CI signal. Cell surface levels of spike not only affected the amplitude of CI but also the kinetics of cell-cell fusion in a concentration dependent manner (**Figure 3B** and **Supplementary Figure 1D**). Although TMPRSS2 has been reported to be an important cellular protease for the activation of S (5, 12, 13), in the context of a cell-cell fusion assay transfection of the A549.ACE2^+^ cells for additional exogenous expression of TMPRSS2 was not a prerequisite to obtain fusion, as also seen by others (25). However, enhanced expression of TMPRSS2 seemed to accelerate and amplify the fusion response (**Figure 3C**).

**FIG 3.**
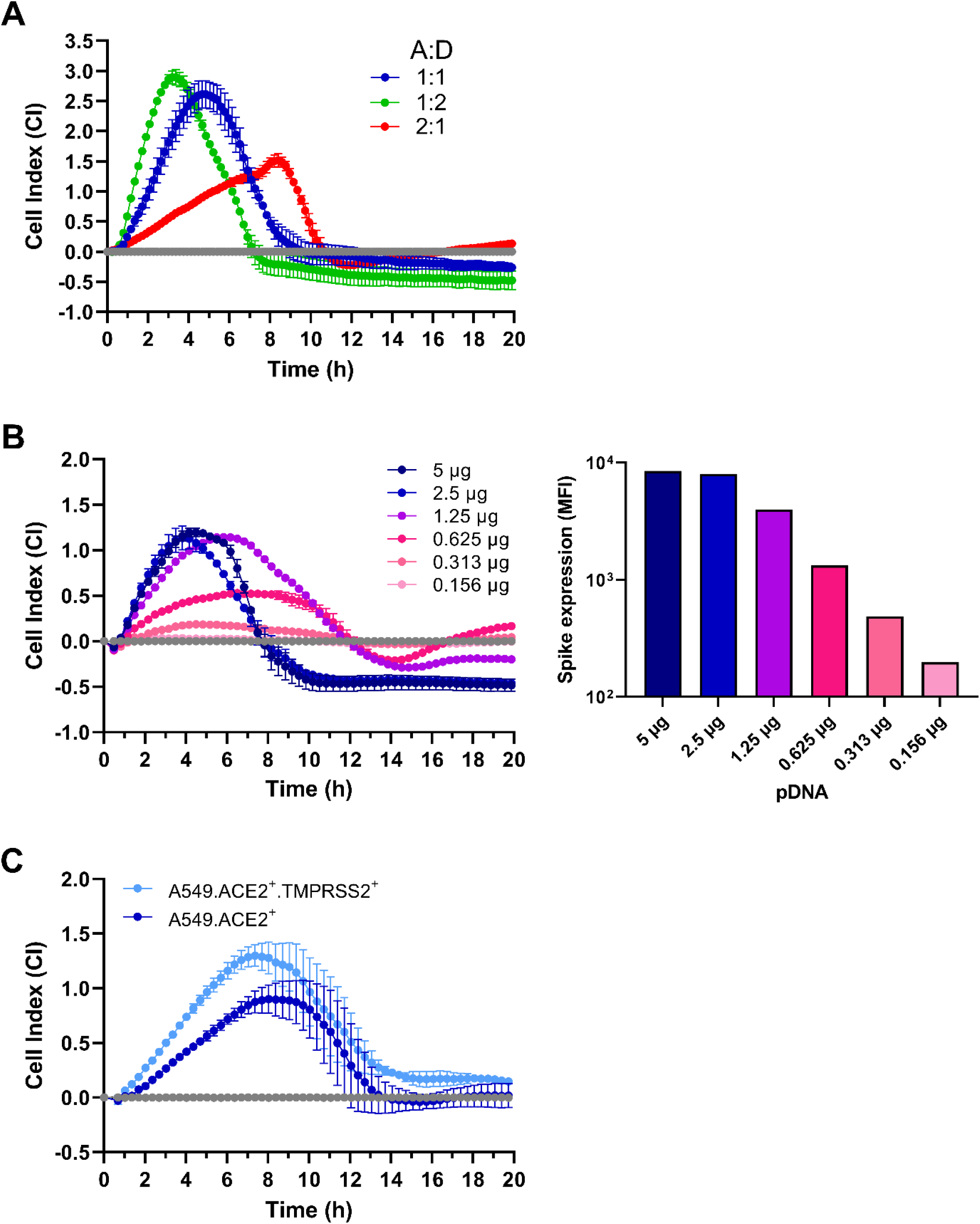
SARS-CoV-2 spike expression correlates with the intensity and kinetics of impedance signal in CEI quantified cell-cell fusion assay. (**A**) Different ratios of A549.ACE2^+^ acceptor (A) and trypsinized Wuhan-Hu-1 spike-transfected HEK293T donor (D) cells. In the 1:1 cell ratio, 15,000 cells of acceptor and donor were used. The graph depicts the impedance signal (expressed as cell index) over time, starting at the moment of cell overlay, of 4 technical replicates (mean ± SD), normalized to the mock-transfected condition (grey horizontal curve). (**B**) Different amounts of Wuhan-Hu-1 SARS-CoV-2 S expressing plasmid DNA (as indicated) were added to 200 μl transfection mixture for the transfection of 400,000 HEK293T donor cells. The next day, cells were trypsinized and added to an A549.ACE2^+^ acceptor cell monolayer. The graph depicts the impedance signal of 2 technical replicates (mean ± SD), normalized to the mock-transfected condition (grey horizontal curve). Bar histograms at the right show the background-substracted mean fluorescence intensity (MFI) values (on a logarithmic scale) for cell surface S staining (Ab R001) of the transfected cells by flow cytometry. See also Supplementary Figure 1D for corresponding flow cytometric histogram plots. (**C**) Comparison of impedance signal of A549.ACE2^+^ cells either mock-transfected (dark blue) versus TMPRSS2-transfected (light blue) and overlayed by trypsinized Wuhan-Hu-1 SARS-CoV-2 S transfected HEK293T cells. The graph depicts the impedance signal of 2 technical replicates (mean ± SD), normalized to the corresponding mock-transfected HEK293T condition (grey horizontal curve).

### Validation of CEI as a quantifiable method of SARS-CoV-2 spike-induced cell-cell fusion

We next explored if our CEI-based fusion assay could be implemented for the evaluation of fusion inhibitors and the profiling of entry inhibitors for SARS-CoV-2. First, we examined the reported SARS-CoV-2 fusion inhibitor EK1 (26-28), a peptide that mimics the Heptad Repeat domain 2 (HR2) of the viral S protein and interferes with the formation of the 6-helix bundle hairpin complex during fusion (**Figure 1A**). Administration of EK1 inhibited cell-cell fusion, with a nearly complete protection at 10 μM concentration (92% reduction in max CI; **Figure 4A**), and in a concentration-dependent manner (IC_50_ value of 0.13 μM; **Supplementary Figure 2A**), thus, validating our CEI-based fusion assay for the analysis of fusion inhibitors. In contrast, the inhibitory effect of a S-neutralizing antibody (Ab) that binds to the RBD of S (Ab R001) was rather limited and depended strongly on the intrinsic efficiency and kinetics of the experimental cell-cell fusion (**Figure 4B** and **Supplementary Figure 2B**). Even a saturating Ab concentration of 10 μg/ml, that completely prevents virus entry of authentic SARS-CoV-2 virus in Vero E6 and Calu-3 cells, as shown earlier (29, 30), reduced the fusion response by only 45% (**Figure 4B**). Furthermore, when Ab R001 was tested in an additional experiment in which cell-cell fusion occurred more efficiently and rapidly because of exogenous TMPRSS2 (as evidenced by the CI response), no reduction in maximum impedance response could be recorded and only a delay in the fusion kinetics was observed (**Supplementary Figure 2B**). Also, when a peptide that represents the RBD of S was tested in our fusion assay, a comparable shift in the CI peak was observed, presumably because of a competition between soluble RBD and cellular-expressed S for the interaction with ACE2 (**Supplementary Figure 2C**; red curve). These data suggest that attachment inhibitors are less potent in the prevention of the fusion event, and indicate that a CEI-based fusion assay might distinguish between fusion inhibitors and attachment inhibitors. Of note, in the absence of S expressing HEK293T cells, CEI detected some cell morphological changes in the A549.ACE2^+^ monolayer because of RBD binding to the ACE2 receptor (**Supplementary Figure 2C**; green curve).

**FIG 4.**
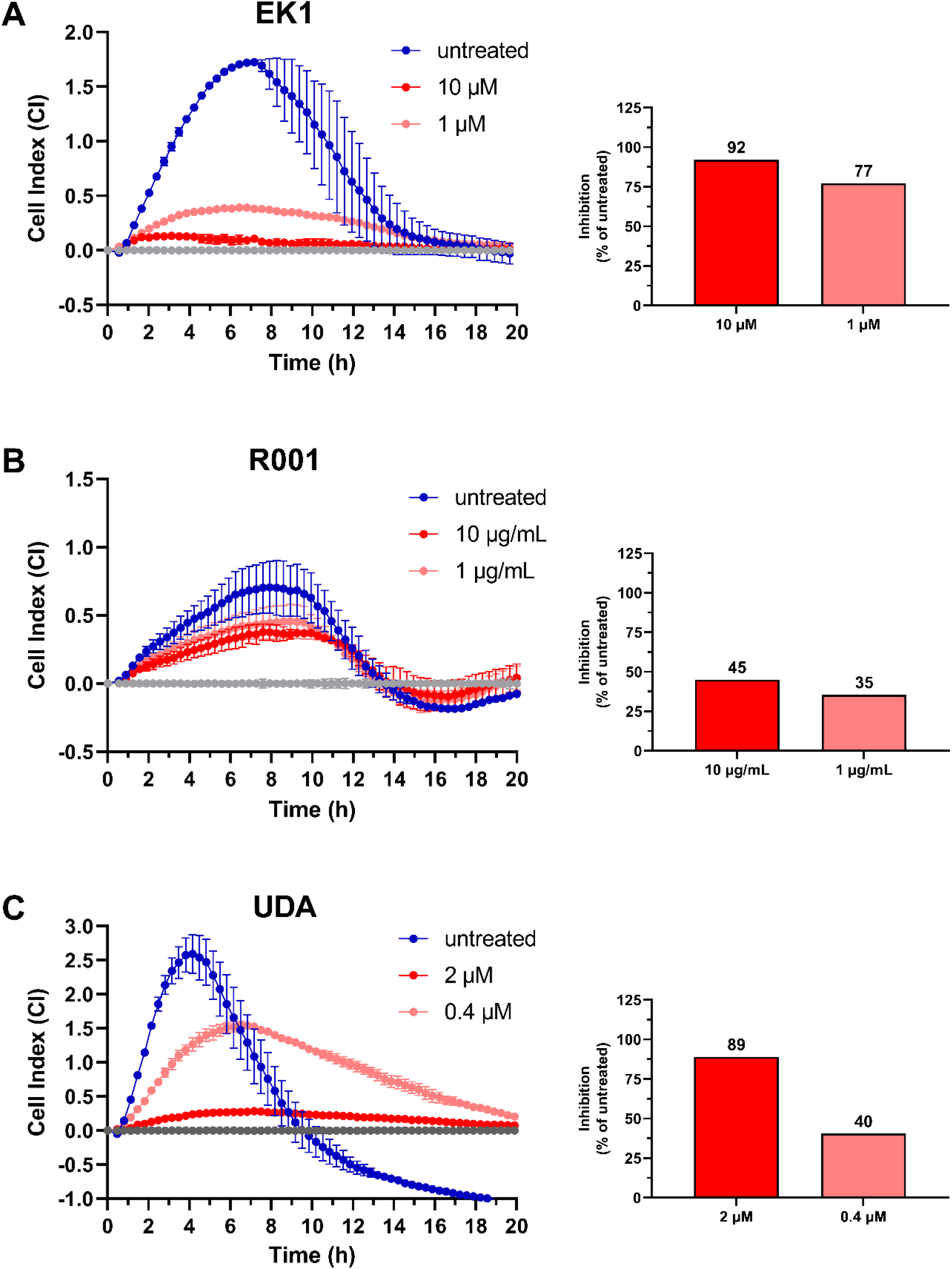
Validation of CEI cell-cell fusion assay with entry inhibitors of SARS-CoV-2. (**A**) The fusion inhibitor EK1 inhibits cell-cell fusion of A549.ACE2^+^ acceptor with S-transfected (20A.EU2 strain) donor cells. Inhibitor and donor cells were added simultaneously to the A549.ACE2^+^ acceptor cells. (**B**) Same as in (A) but for the attachment inhibitor R001, an RBD binding antibody that neutralizes viral entry of authentic SARS-CoV-2 virus, and with SARS-CoV-2 Wuhan-Hu-1 S. (**C**) Same as in (A) but for the entry inhibitor UDA, a carbohydrate-binding small monomeric plant lectin from stinging nettle rhizomes. The graphs on the left depict the impedance signal of 2 technical replicates (mean ± SD), normalized to the mock-transfected condition (grey horizontal curve). Bar histograms on the right show the inhibition of impedance response relative to the untreated control sample, calculated from the maximum CI values obtained for each treated sample.

In a previous study, we investigated the antiviral potential of the plant lectin UDA against SARS-CoV-2 and reported a profound inhibitory effect of UDA on virus entry (23). Evaluation of UDA in our CEI assay clearly demonstrated deceleration and inhibition of S-induced fusion in a concentration-dependent way (**Figure 4C**), with an IC_50_ value of 0.55 μM, that correlates well with its reported antiviral potency against authentic SARS-CoV-2 *in vitro* (23). In line with that report (23), UDA pretreatment of the ACE2^+^ acceptor cells (followed by compound wash-out) had little impact on cell-cell fusion whereas pre-incubation of the S-expressing cells with UDA (followed by compound wash-out) profoundly protected the fusion event (**Supplementary Figure 3**), confirming our observation that UDA binds to the glycosylated viral S protein rather than the cellular receptor.

### CEI to analyze the fusogenic potential of SARS-CoV-2 spike variants

One of the advantages of our CEI assay is the flexibility and easy-to-adapt format of the protocol. For example, mutants of SARS-CoV-2 spike can be easily designed and used for the transfection of the HEK293T cells. Hence, with CEI spikes from different SARS-CoV-2 VOCs can be compared for their fusogenic potential. As shown in **Figure 5A**, spikes carrying the D614G mutation, as present in all variants since early 2020 (Nextstrain clade 20A and its descendants), retained a similar fusogenic efficiency as compared to the original Wuhan-Hu-1 strain S. In contrast, the Omicron VOC circulating in 2022 has been reported to possess a reduced fusogenic potential (9, 10, 31). Interestingly, our CEI assay also confirmed the reduced fusion of Omicron S (BA.1 variant) as compared to Wuhan-Hu-1 S (**Figure 5B**), as evidenced by a slower and attenuated impedance response. This reduced fusogenic potential of Omicron was also observed when an excess of spike-expressing plasmid was used for the transfection of the HEK293T cells (**Supplementary Figure 4A**). In fact, the reduced fusion potential of Omicron as compared to Wuhan-Hu-1 could not directly be linked to attenuated spike expression, as evidenced by the comparable levels of S protein that could be detected by flow cytometry on the surface of HEK293T cells transfected with similar amounts (2.5 μg) of plasmid (**Supplementary Figure 4B**).

**FIG 5.**
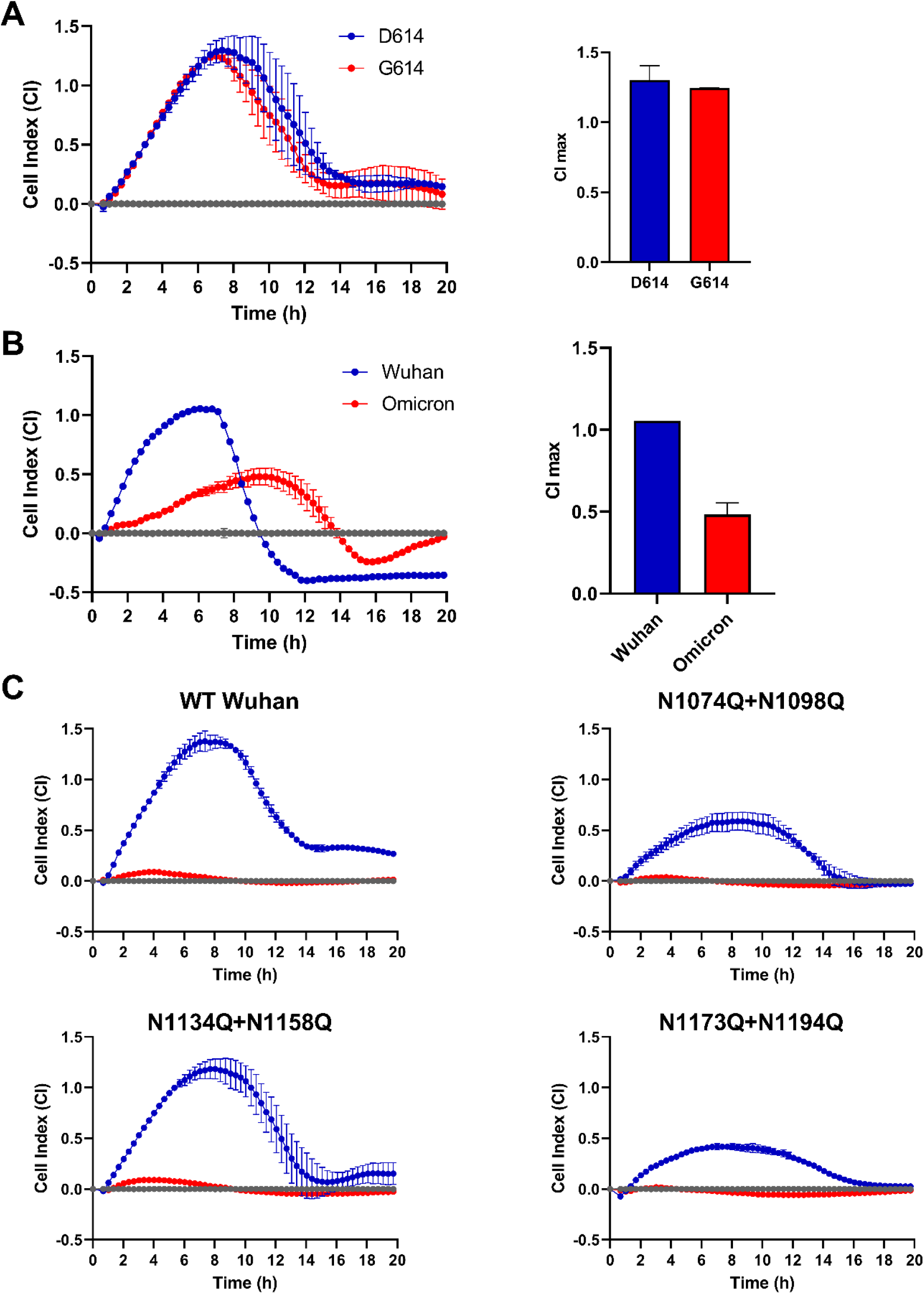
CEI measures the alteration in fusogenic potential of SARS-CoV-2 S variants. (**A**) Comparison of impedance signal of A549.ACE2^+^ cells overlayed with HEK293T cells transfected with plasmid DNA coding for SARS-CoV-2 S either from Wuhan-Hu-1, carrying D614 (blue) or a mutant with G614 (red) as found in the Nextstrain clade 20A and its descendants. In both conditions, 2.5 μg S expressing plasmid DNA was added to 200 μl transfection mixture for the transfection of 400,000 HEK293T donor cells. Graph on the left depicts the impedance signal of 2 technical replicates (mean ± SD), normalized to the mock-transfected condition (grey horizontal curve). Bar histograms on the right show the maximum CI values (mean ± SD). (**B**) Same as in (A) but for the comparison between Wuhan-Hu-1 and Omicron. (**C**) Fusion-inhibitory effect of UDA (2 μM) on different N-glycosylation deletion mutants. Mutants of Wuhan-Hu-1 S that contained two deletions of adjacent N-glycosylation sites in the S2 subunit were generated (by N into Q conversion) and analyzed in a CEI-based cell-cell fusion assay for their sensitivity to UDA. Graphs show the impedance signal of 2 technical replicates (mean ± SD), normalized to the mock-transfected condition.

Finally, we used CEI to analyze different SARS-CoV-2 spike glycosylation deletions to complement previous work on the plant lectin UDA (23). As shown in **Figure 5C**, a fusion assay with 3 different double-glycosylation mutants confirmed the preserved activity of UDA against spike variants with depleted glycans in the S2 unit, indicating that removal of N-glycosylation sites in the S2 subunit will not result in SARS-CoV-2 escape mutants for UDA. Of note, the difference in the CI amplitude that was observed between the untreated S-transfected control samples of the different N-glycosylation mutants (**Supplementary Figure 4C**) was simply related to different levels of S expression on the HEK293T cells, as verified by flow cytometry (**Supplementary Figure 4D**), presumably because N-glycosylation depletion affected S protein stability and subsequent cell surface expression.

## Discussion

In this study, we have implemented cell-based electrical impedance to measure SARS-CoV-2 spike-induced cell-cell fusion in real time and have demonstrated that CEI can be used to characterize fusion inhibitors for SARS-CoV-2. In addition, we have shown its feasibility to analyze the fusogenic properties of spike proteins from different circulating SARS-CoV-2 VOCs. The technique measures the kinetics of the fusion process in real-time, in a label-free and in a non-invasive format, allowing further down-stream analysis of the samples (e.g., qPCR or Western blot analysis). Although CEI does not require expensive detection reagents or artificial detection systems (32), specialized microtiter plates embedded with gold electrodes are needed for the experimental set-up, which may affect the cost of this alternative fusion assay. A big advantage of CEI as compared with other cell-cell fusion detection methods for SARS-CoV-2 (e.g., split neongreen; **Figure 1B**) is the real-time quantification of membrane fusion, which does not depend on the intermingling of the cytosolic content of the fused cells and the formation of an active fluorescent protein or translation of a reporter enzyme (e.g., luciferase (25)). Generally, S induced cell-cell fusion already manifests within the first hours of cell overlay (**Figure 1C**; syncytia visible at 3h), but needs several more hours of incubation to obtain a quantifiable reporter signal (**Figure 1C**; fluorescent signal at 12h). In contrast, with CEI even a small delay or minor reduction in the cell-cell fusion process can be monitored for inhibitors or mutant spike proteins.

Routine antiviral screens often rely on phenotypic assays, such as the reduction in virus-induced cytopathic effect (CPE) or plaque formation, and generally make use of a microscopic, luminescent, fluorescent, or colorimetric readout. Some of the disadvantages of these techniques are that can be slow (multiple days), are typically time consuming, require multiple handling steps and rely on optimal endpoint selection in order to achieve good assay quality. CEI offers a valuable addition to this array of antiviral methods with real-time measurements and an objective readout. Furthermore, CEI is able to measure nanoscale morphological changes and is thus a more sensitive tool to analyze cellular processes such as highly dynamic membrane fusion. Also, a CEI-based fusion assay can be performed with fusion protein-expressing cells as an alternative to authentic virus, allowing antiviral analysis of entry inhibitors at a lower biosafety level, which might be of particular interest for pathogens that require BSL3 facilities.

Entry of viruses into host target cells is an attractive target for antiviral intervention (33). Inhibiting the virus before it enters the host cell, e.g., by targeting the viral fusion machinery, is a potent antiviral strategy that has been successful for the treatment of human immunodeficiency virus (34). Furthermore, as these membrane fusion processes are critical for viral infection, targets are often well conserved across different viral families (35, 36), suggesting the potential for developing pan-viral broad spectrum inhibitors.

Here, we exploited CEI to analyze the activity of fusion inhibitors. In our comparative study of entry inhibitors for SARS-CoV-2, we observed differential potency in our CEI-based cell-cell fusion assay between an attachment inhibitor (Ab R001) and a fusion inhibitor (EK1). The limited fusion inhibitory effect of the S-binding R001 might be related to the excessive expression of the spike protein on the surface of transfected HEK293T cells, which greatly differs from the number of fusion proteins on the viral envelop in antiviral CPE-based cellular assays. In that perspective the concentration Ab (10 μg/ml) used in our cell-cell fusion experiments was most likely suboptimal. Accordingly, Zhao *et al*. observed for anti-spike monoclonal antibodies 15-20 fold less activity against cell-cell fusion as compared to pseudotyped SARS-CoV-2 virus infection (37). In addition, in a recent reported luciferase-based reporter gene fusion assay for SARS-CoV-2, a concentration of 28 μg/ml Ab R001 resulted in approximately 60-70% inhibition of cell-cell fusion, thus, still no complete protection (25), however, the spike-expressing cells were pre-incubated with the Ab before the overlay. Thus, it seems that attachment inhibitors have less opportunity to prevent the initiation of the fusion process once the spike proteins are being triggered by ACE2. Nevertheless, antivirals that interfere with the conformational changes and refolding of the S2 subunit have a stronger inhibitory effect on membrane fusion and can be clearly identified through CEI. Our data also indicated that expression of TMPRSS2 is not a prerequisite to obtain S-mediated cell-cell fusion. In a CEI experimental setting, other host proteases, such as matrix metalloproteases might activate the SARS-Cov-2 S glycoprotein during the process of syncytium formation (38).

Interestingly, CEI can also be employed to study the fusogenic properties of spike proteins from different SARS-CoV-2 VOCs. In line with several reports (9, 10, 31), with our CEI based cell-cell fusion assay we also measured a reduced fusion activity for the spike protein of the Omicron BA.1 variant. Although subsequent flow cytometric analysis of the spike levels on the surface of the transfected HEK293T cells did not indicate an attenuated protein expression for Omicron (as compared to the original Wuhan-Hu-1), flow cytometry does not provide details on the amount of processed (S1/S2 cleaved) spike or the trimeric nature of the SARS-CoV-2 fusion protein. As we demonstrated that the cell-cell fusion process is strongly depending on the amount of wild-type spike protein expressed on the surface of the HEK293T cells, the proteolytic state of S will undoubtedbly contribute to the efficiency of the fusion process.

In summary, we have developed an assay that makes use of cell-based electrical impedance to monitor in real time cell-cell fusion for SARS-CoV-2, which provides a powerful tool to investigate specific cell membrane fusion events. The CEI technique can add valuable information on the fusion process of circulating enveloped viruses and on the mode of action of new antivirals. As such, CEI can support profiling efforts of novel potent fusion inhibitors for SARS-CoV-2, and in turn can help in the development of next generation inhibitors with improved antiviral activity.

## Materials and Methods

### Cell lines

Human Embryonic Kidney 293T (HEK293T) cells (Cat. No. CRL-3216) and Human adenocarcinomic alveolar epithelial cells A549 (Cat. No. CCL-185), were obtained from ATCC as mycoplasma-free stocks and grown in Dulbecco’s Modified Eagle Medium (DMEM, Thermo Fisher Scientific) supplemented with 10% (*v* / *v*) fetal bovine serum (FBS; HyClone). A549.ACE2^+^ cells were generated by stably transducing A549 with ACE2 (23), by second generation lentiviral transduction with a lentiviral transfer vector containing the human ACE2 coding sequence as described elsewhere (24). Cell lines were maintained at 37°C in a humidified environment with 5% CO_2_. Cells were passaged every 3 to 4 days.

### Antibodies and compounds

#### Antibodies

The following antibodies were used for flow cytometry: rabbit monoclonal SARS-CoV-2 spike-specific antibody [R001] (Cat. n° 40592-R001, Sino Biological), mouse monoclonal SARS-CoV-2 spike-specific antibody [MM57] (cat. n° 40592-MM57, Sino Biological), fluorescently-labelled Alexa Fluor 647 (AF647) goat anti-Rabbit IgG monoclonal antibody (Cat. n° 4414, Cell Signaling Technologies), and phycoerythrin (PE)-labelled goat anti-mouse IgG (cat. n° 405307, BioLegend).

#### Compounds

Wuhan-Hu-1 SARS-CoV-2 receptor binding domain was purchased from Sino Biological (2019-nCoV spike RBD, Cat n° 40592-VNAH). Urtica dioica agglutinin (UDA) from Stinging Nettle was obtained from EY Laboratories, CA, USA (Cat. n° L-8005-1). EK1, with amino acid sequence GSLDQINVTFLDLEYEMKKLEEAIKKLEESYIDLKELG, was synthesized as a custom peptide with N-terminal acetylation and C-terminal amidation (Life Technologies Europe Bv).

### Plasmids

All plasmids were generated with the NEBuilder DNA assembly kit (New England Biolabs), using a pCAGGS plasmid digested with EcoRV-HF and HindIII-HF (New England Biolabs) as backbone. For pCAGGS.SARS-CoV-2_SΔ19 a PCR fragment encoding codon-optimized SARS-CoV-2 Wuhan-Hu-1 spike protein (amplified from pCMV3-C-Myc; VG40589-CM, SinoBiological) with a C-terminal 19 amino acid deletion as described in (39) was cloned into the pCAGGS backbone. For pCAGGS.SARS-CoV-2_SΔ19_fpl_mNG2(11) the same cloning strategy was used as the previous plasmid, but with the addition of a PCR fragment containing a 12 amino acid flexible protein linker (fpl) and a modified 11^th^ betasheet of mNeonGreen (mNG2(11)) (40). For pCAGGS.SARS-CoV-2 SΔ19 [D614G], PCR was performed on the codon-optimized SARS-CoV-2 Wuhan spike sequence in two parts to introduce a D614G amino acid mutation in the overlapping region between fragments. For the pCAGGS.SARS-CoV-2_SΔ19 [N1074Q+N1098Q]_fpl_mNG2(11), pCAGGS.SARS-CoV-2_SΔ19 [N1134Q+N1158Q]_fpl_mNG2(11) and pCAGGS.SARS-CoV-2_SΔ19 [N1173Q+N1194Q]_fpl_mNG2(11) mutant plasmids, mutations were again introduced in overlaps of the PCR fragments obtained from PCR of the codon-optimized Wuhan spike sequence. Fragments were ligated in combination with a PCR fragment containing the fpl and mNG2(11) sequences. pCAGGS.SARS-CoV-2_S Δ19_Omicron_fpl_mNG2(11) was cloned via PCR performed on cDNA synthesized from an RNA extract of a SARS-CoV-2 Omicron BA.1 virus stock (SARS-CoV-2 B.1.1529). Fragments were again ligated with a PCR fragment containing the fpl and mNG2(11) sequences. pCAGGS.SARS-CoV-2_S Δ19_Omicron-Opt_fpl_mNG2(11) was assembled from a PCR performed on a codon-optimized SARS-CoV-2 Omicron BA.1 sequence synthesized by Genscript. Fragments were assembled with a PCR fragment containing the fpl and mNG2(11) sequences. pcDNA3.1.mNG2(1-10) was generated through NEBuilder DNA assembly of a pcDNA3.1 vector (Thermo Fisher Scientific), amplified by PCR, and 10 betasheets of a modified mNeonGreen synthesized by Genscript. For the transfection of TMPRSS2, the pcDNA3.1+ plasmid encoding TMPRSS2-DYK was purchased from Genscript. All plasmids were sequence-verified before use with Sanger sequencing (Macrogen).

### Transient transfection

Prior to transfection HEK293T and A549 cells were plated in 6-well pates to reach 50-70% and 80-90% confluency, respectively, after an overnight incubation at 37 °C. Lipofectamine LTX (Invitrogen) was used for the transfection of plasmid DNA according to the manufacturer’s protocol.

### Split neongreen cell-cell fusion assay

Transfection mixes were prepared with 2.5 μg pCAGGS.SARS-CoV-2_SΔ19_fpl_mNG2(11)_opt plasmid encoding for SARS-CoV-2 spike protein for HEK293T transfection; and 2.5 μg pcDNA3.1.mNG2(1-10) for A549.ACE2^+^ transfection. HEK293T cells were allowed to incubate for 24 h for efficient exogenous spike protein expression. At 6 h post transfection, transfected A549.ACE2^+^ cells were digested with 0.05% trypsin, washed, resuspended and counted on a Luna cell counter (Logos Biosystems), added to a 96-well plate at 2.2 × 10^4^ cells per well and incubated for 18 h. Next day, transfected HEK293T cells were collected, digested with 0.25% trypsin, washed, resuspended, counted and administered to the A549.ACE2^+^ cells at 2 × 10^4^ cells per well. Fusion events were visualized for 24 h at 20 min intervals using the IncuCyte® S3 Live-Cell Analysis System (Sartorius). Phase contrast and GFP images (4 per well) were taken using a 20x objective lens at 10-minute intervals for a 5 hours period, and 1 hour intervals afterwards. Image processing was performed using the IncuCyte® software.

### Cellular Electrical Impedance cell-cell fusion assay

The xCELLigence Real-Time Cell Analyzer (RTCA) DP instrument (Agilent, Santa Clara, CA, USA) was used to measure changes in cellular impedance following addition of cells on top of the monolayer. Briefly, RTCA E-plate VIEW 16 plates with embedded golden electrodes (#300600880, Agilent, Santa Clara, CA, USA) were used for the experiments. First a blank measurement of a sensor E-Plate VIEW 16 PET was performed (only medium). This was followed by the addition of 15,000 A549.ACE2^+^ cells (in growth medium supplemented with 2% FBS) to each well. E-plates were placed at room temperature for 15 min and then transferred to the xCELLigence RTCA instrument, located in an incubator at 37 °C and 5% CO2. The attachment and overnight growth of the cells was monitored (measurement every 20 min). Cell adherence and growth result in an increase in CI followed by a flattening of the curve when the cells reach confluency. In parallel, HEK293T cells (400,000 cells per well in growth medium supplemented with 10% FBS) were transfected with an S-expressing plasmid. Following overnight incubation, a short CEI normalisation measurement (5 consecutive measurements, every 5 s) was performed on the A549 cell monolayer. In parallel, spike-expressing transfected HEK293T cells were collected, digested with 0.25% trypsin, washed, resuspended, counted and administered to the A549.ACE2^+^ cells at 15,000 cells per well (= overlay step), simultaneously with the test compounds or vehicle control. To quantify spike-independent CI changes resulting from the overlay, an equal number of HEK293T cells mock-transfected with empty vector is added to the A549.ACE2^+^ cells instead. After the overlay, the A549.ACE2^+^ monolayer is monitored over time for 24h and data points are displayed every 2 minutes.

### CEI data analysis

The CEI biosensor monitors the Cell Index (CI), a dimensionless parameter derived from the frequency-dependent resistance (R) component of the impedance value (Z) at 10, 25 and 50 kHz frequency. Raw CI values were used as a starting point for data manipulations. All data are first normalized to the baseline before the overlay step, to reduce inter-well variation. Spike-dependent fusion was calculated by subtracting the CI values of A549.ACE2^+^:HEK293T.empty_vector overlay (spike independent) from the CI changes of the A549.ACE2^+^:HEK293T.spike overlay (spike-dependent + independent). This results in a baseline-corrected normalized CI measure. The maximal CI change of the A549.ACE2^+^:HEK293T.spike overlay (of the baseline corrected CI value) in the absence of compound, is then set to 100% and the maximal CI change of the conditions with compound are reported relative to this value. CEI data were preprocessed, normalized and baseline-corrected using an in-house built Matlab script (version R2016b, Mathworks). IC_50_ calculation was done in GraphPad Prism (version 9) using nonlinear regression: log[inhibitor] vs. normalized response variable slope.

### Flow cytometry

For the cell surface staining of spike-transfected HEK293T cells, cells were collected, washed in PBS, resuspended, transferred to tubes and samples were centrifuged in a cooled centrifuge (4 °C) at 500 *g* for 5 min. After removal of the supernatant, cells were incubated with the primary (anti-spike) antibody (30 min at 4 °C), washed in PBS, followed by a 30 min incubation at 4 °C with the secondary (labeled) antibody, and washed again. Finally, samples were stored in PBS containing 2 % formaldehyde (VWR Life Science AMRESCO). Acquisition of all samples was done on a BD FACSCelesta flow cytometer (BD Biosciences) with BD FACSDiva v8.0.1 software. Flow cytometric data were analyzed in FlowJo v10.1 (Tree Star). Subsequent analysis with appropriate cell gating was performed to exclude cell debris and doublet cells, in order to acquire data on living, single cells only.

### Statistical analysis

Data were visualized as means ± standard deviation (SD) and were analyzed using GraphPad Prism 9.3.1 software.

## Abbreviations

ACE2: angiotensin-converting enzyme 2
CEI: cellular electrical impedance
CI: cell index
COVID-19: coronavirus disease 2019
CTS: cathepsin
FP: fusion peptide
MFI: mean fluorescence intensity
RBD: receptor-binding domain
SARS-CoV-2: severe acute respiratory syndrome coronavirus 2
S: spike
TMPRSS2: transmembrane serine protease 2

## Supplemental material

Supplemental material is available online only.

**Supplemental file 1**, PDF file, 0.97 MB

Supplemental movie 1, MP4 file, 19.6 MB

Suppelemental movie 2, MP4 file, 17.5 MB

## Acknowledgements

We thank Geert Schoofs for technical support to the flow cytometry experiments, and Anita Camps and Eef Meyen for their help with the cell cultures. Images were created with support from BioRender.com.

## Author contributions

K.V., E.V., J.D. and J.S. designed research; E.V., J.D., E.M., B.P., J.S. and S.N. performed research; E.V., J.D., J.S., S.N. and K.V. analyzed the data; E.V., J.D., A.S. and K.V. wrote the manuscript; D.S. contributed new reagents/analytic tools. All of the authors discussed the results and commented on the manuscript.

## SUPPLEMENTAL MATERIAL

### Supplementary Figures

**Suppl FIG 1.**
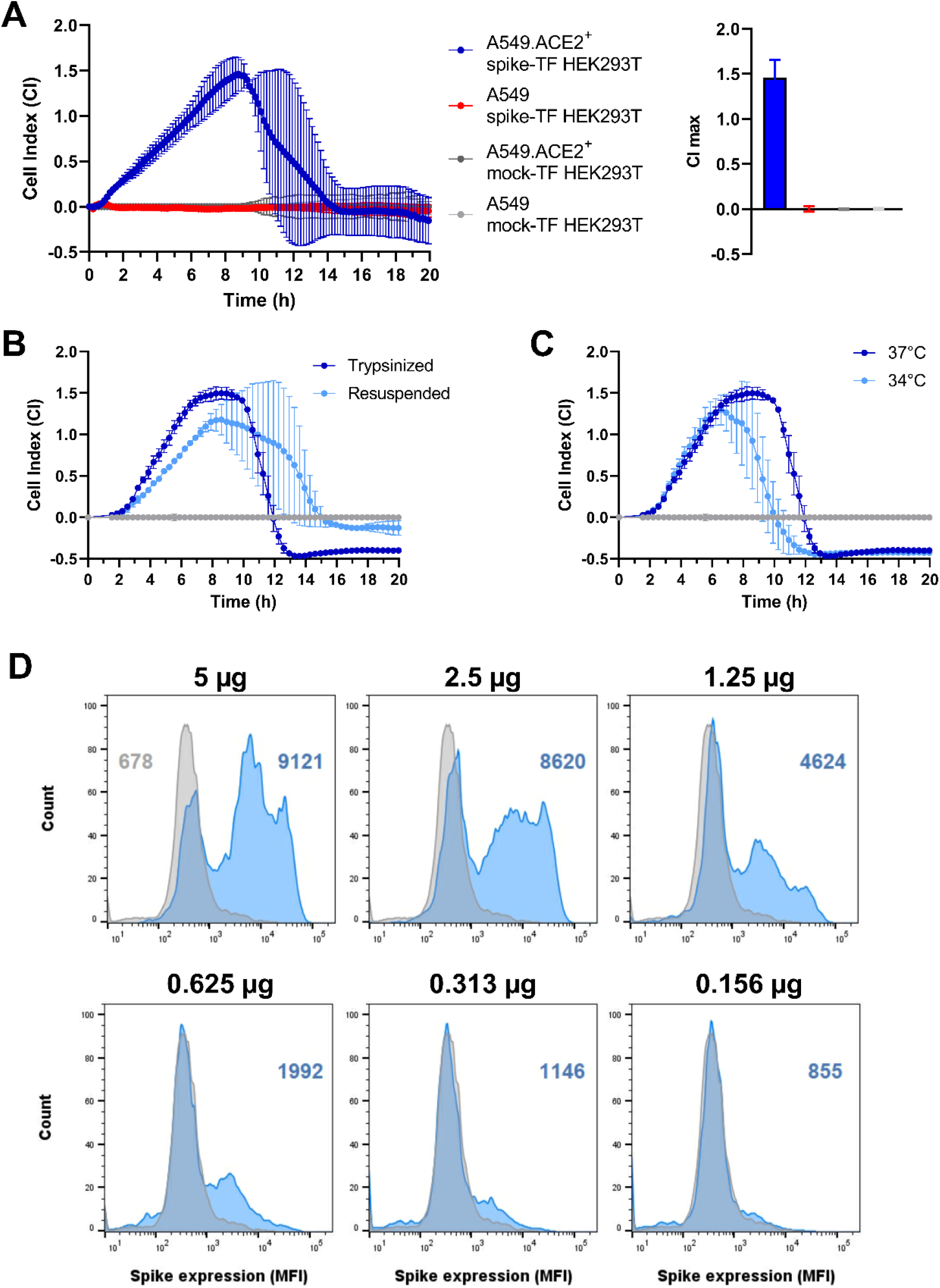
Optimization of CEI-measured S-induced cell-cell fusion. (**A**) Cell-cell fusion depends both on the expression of ACE2 on A549 cells and SARS-CoV-2 S protein on transfected HEK293T cells. Different combinations of acceptor and donor cells were tested as indicated. The graph depicts the impedance signal of 4 technical replicates (mean ± SD). Bar histograms at the right represent the cell index value at 8:43h post overlay, when the maximum was reached in the positive control. Note that no increase in impedance signal (CI max ∼ 0) was obtained in the conditions in which ACE2 and/or spike were not (over)-expressed. (**B**) Comparison of impedance signal of A549.ACE2^+^ cells overlayed with SARS-CoV-2 Wuhan Hu-1 S-transfected HEK293T cells, either trypsinized or collected by resuspending. The graph depicts the impedance signal of 2 technical replicates (mean ± SD). (**C**) HEK293T cells were transfected with SARS-CoV-2 Wuhan Hu-1 S. After 6h, transfection reagent was removed and cells were incubated either at 34°C or 37°C for 18h. S-expressing cells were then trypsinized, collected and administered to a A549.ACE2^+^ cell monolayer, and further incubated at 37°C for the CEI measurement. The graph depicts the impedance signal of 2 technical replicates (mean ± SD). (**D**) Flow cytometric histogram plots of the samples presented in Fig 3B (see figure legend to Fig 3B for experimental details). HEK293T cells were collected 24h post transfection, stained with anti-S Ab (R001) and an AF647-labeled secondary Ab. Of each sample 10,000 cells were analyzed on a FACSCelesta to calculate the mean fluorescence intensity (MFI) value. The grey histogram represents the stained mock-transfected background control sample, whereas the S-transfected cells are indicated in blue. The values in color refer to the respective MFI value.

**Suppl FIG 2.**
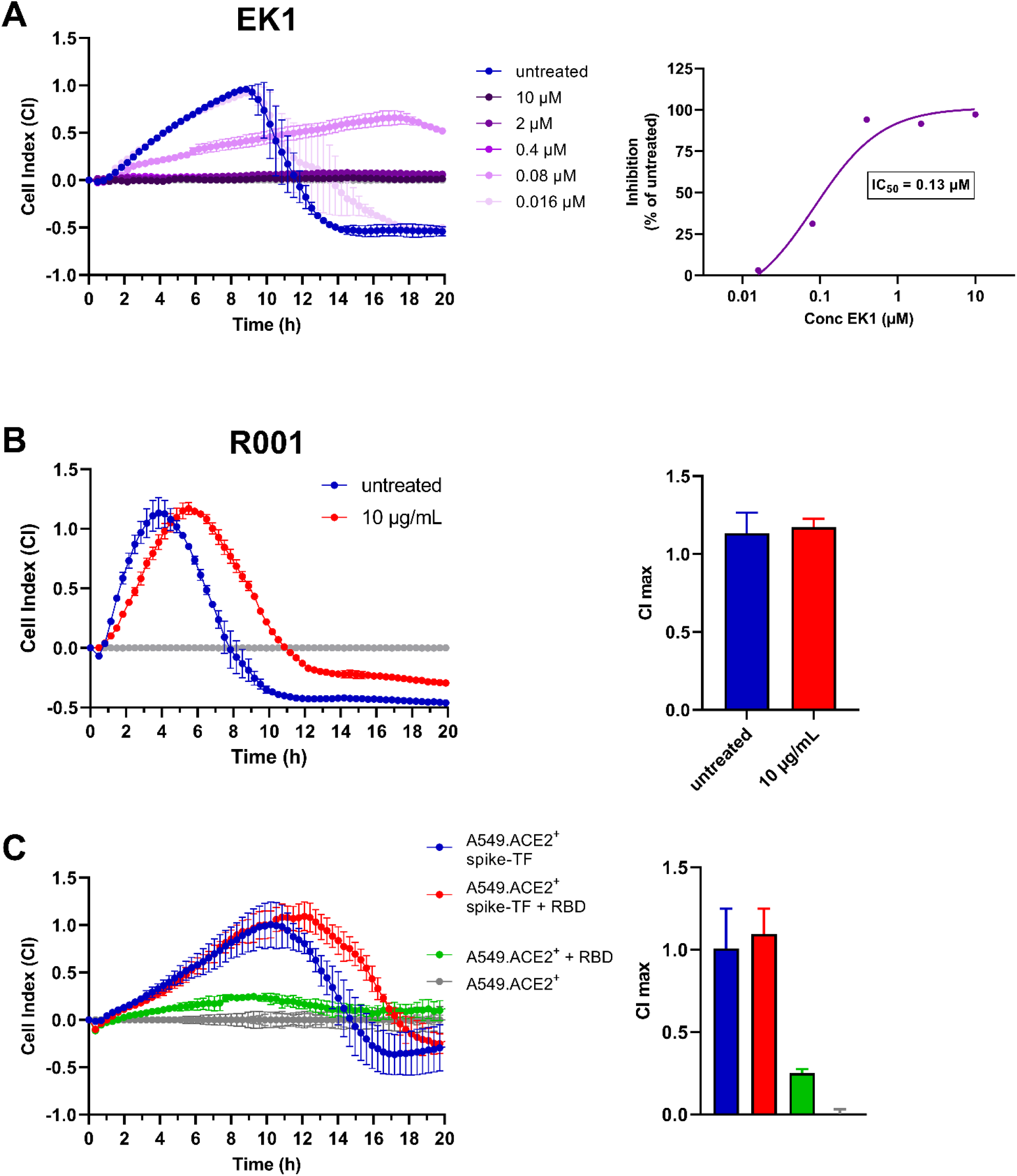
Validation of CEI cell-cell fusion assay with entry inhibitors of SARS-CoV-2. (**A**) Fusion inhibitor EK1 inhibits concentration-dependently the cell-cell fusion of S-transfected (Wuhan-Hu-1 strain) HEK293T donor with A549.ACE2^+^ acceptor cells. Inhibitor and donor cells were added simultaneously to the A549.ACE2^+^ acceptor cells. The graph on the left shows the impedance signal of 2 technical replicates (mean ± SD), normalized to the mock-transfected condition. Concentration-response curve on the right shows the inhibition of impedance response relative to the untreated control sample, calculated from the CI values obtained at the time point when maximum CI was reached in the positive control. The calculated 50% inhibitory concentration (IC_50_) is given in the boxed insert. (**B**) The S-binding attachment inhibitor Ab R001 delays the the cell-cell fusion of S-transfected (Wuhan-Hu-1 strain) HEK293T donor with A549.ACE2^+^ acceptor cells transiently transfected with TMPRSS2. R001 (10 μg/ml) and donor cells were added simultaneously to the A549.ACE2^+^.TMPRSS2^+^ acceptor cells. The graph shows the impedance signal of 2 technical replicates (mean ± SD), normalized to the mock-transfected condition. Bar histograms on the right show the maximum CI values (mean ± SD; n=2). (**C**) RBD peptide from SARS-CoV-2 Wuhan-Hu-1 S delays the cell-cell fusion of S-transfected (Wuhan-Hu-1) HEK293T donor with A549.ACE2^+^ acceptor cells. RBD (81 nM) and donor cells were added simultaneously to the A549.ACE2^+^ acceptor cells (red curve). In parallel, RBD (81 nM) was administered to a monolayer of A549.ACE2^+^ cells in the absence of spike-expressing cells to measure the (small) morphological changes induced by RBD binding to the ACE2 receptor (green curve). The graph shows the impedance signal of 4 technical replicates (mean ± SD), normalized to the respective mock-transfected or untreated condition. Bar histograms on the right show the maximum CI values (mean ± SD; n=4).

**Suppl FIG 3.**
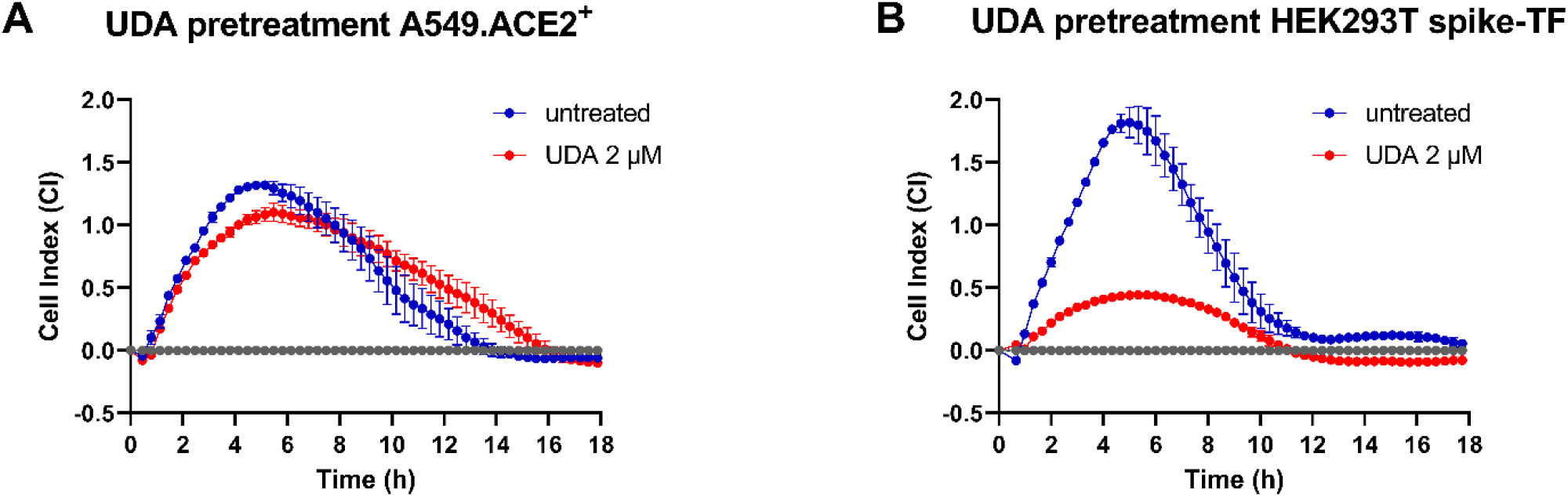
Plant lectin UDA inhibits CEI quantified cell-cell fusion through binding to SARS-CoV-2 S. (**A**) A monolayer of A549.ACE2^+^ cells were pretreated with UDA (2μM) for 1h at 37°C, washed and overlaid with S-transfected (Wuhan-Hu-1) HEK293T cells without additional compound administration. (**B**) At 24h post transfection, S-transfected (Wuhan-Hu-1) HEK293T cells were first pretreated with UDA (2μM) for 1h at 4°C, trypsinized, collected and washed. Cells were resuspended in culture medium and overlaid on a monolayer of A549.ACE2^+^ cells without additional compound administration. Graph show the impedance signal of 2 technical replicates (mean ± SD), normalized to the mock-transfected condition.

**Suppl FIG 4.**
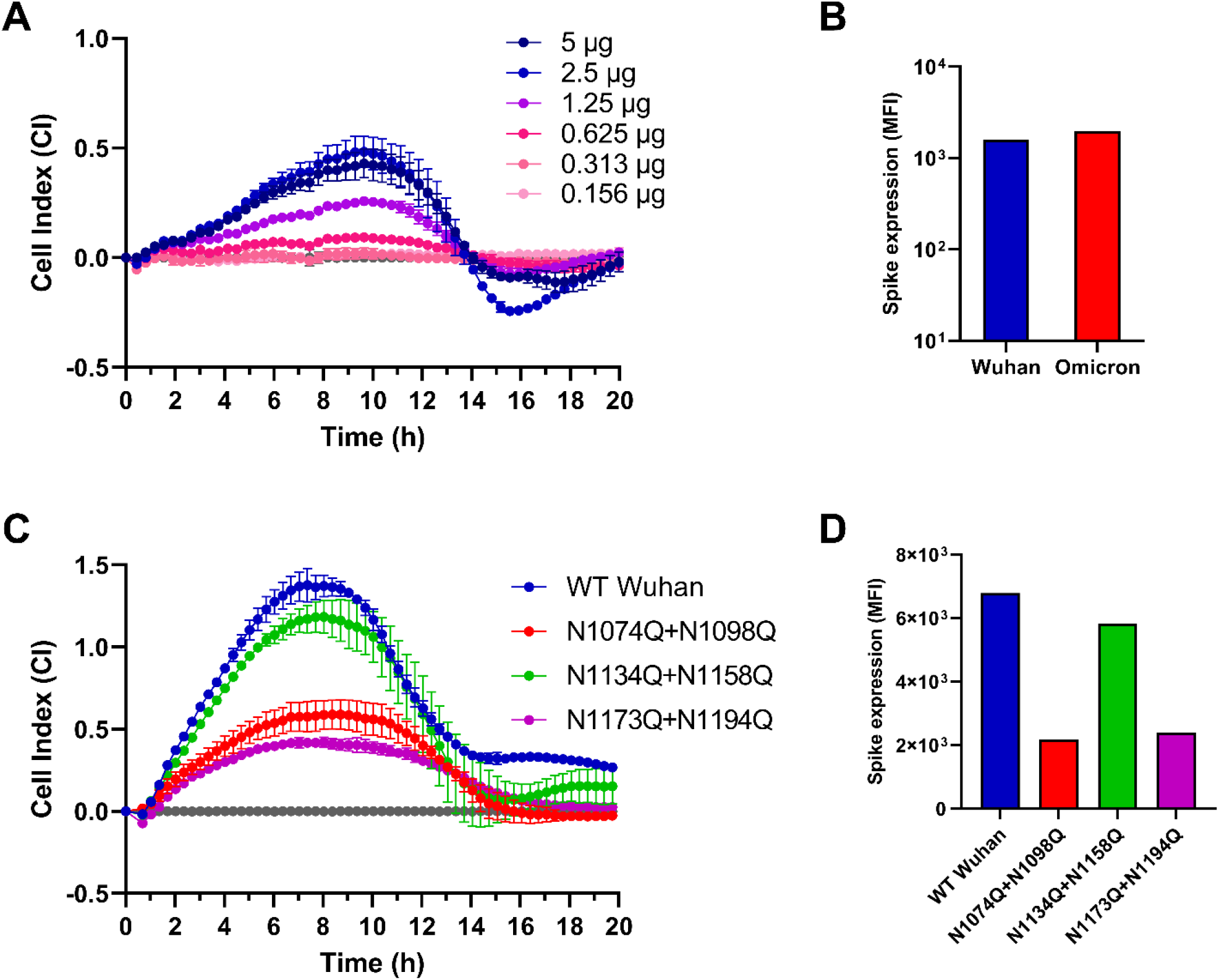
CEI measures the alteration in fusogenic potential of SARS-CoV-2 S variants. (**A**) Same as in Fig 3B but with transfection of Omicron S (BA.1 variant). (**B**) Transfected HEK293T samples (each with 2.5 μg plasmid DNA) from Fig 5B were collected 24h post transfection, stained with anti-S Ab (R001) and an AF647-labeled secondary Ab. Bar histograms represent the background-corrected mean fluorescence intensity (MFI) values (on a logarithmic scale), calculated from 10,000 cells analyzed by a FACSCelesta flow cytometer. (**C**) Untreated control samples from Fig 5C were plotted together in one graph to compare the fusion efficiency of N-glycosylation mutants of S. (**D**) Transfected HEK293T samples from (C) were collected 24h post transfection, stained with anti-S Ab (MM57) and an PE-labeled secondary Ab. Bar histograms represent the background-corrected MFI values (on a logarithmic scale), calculated from 10,000 cells analyzed by a FACSCelesta flow cytometer.

### Supplementary movies

**Supplementary movie 1**. A549.ACE2^+^ cells (transfected to express the first 10 betasheets of neongreen) were overlayed with HEK293T cells co-transfected with a plasmid encoding the SARS-CoV-2 spike protein and a plasmid encoding the 11^th^ betasheet of neongreen. Overlay was done with mock-transfected (empty vector; left) or spike-transfected (middle and right) HEK293T cells, either in the absence (middle) or presence (right) of the fusion inhibitor (FI) EK1 (2 μM). Fusion events were visualized using the IncuCyte® S3 Live-Cell Analysis System (Sartorius). Phase contrast and GFP images were taken using a 20x objective lens at 30 minute intervals for a 24 hours period. Image processing was performed using the IncuCyte software.

**Supplementary movie 2**. A549.ACE2^+^ cells were overlayed with HEK293T cells transfected with a plasmid encoding the Wuhan-Hu-1 SARS-CoV-2 spike protein. Overlay was done with mock-transfected (empty vector; red) or spike-transfected (blue) HEK293T cells. Left panels: fusion events were visualized using the IncuCyte® S3 Live-Cell Analysis System (Sartorius). Phase contrast images were taken using a 20x objective lens at 30 minute intervals for a 19 hours period. Image processing was performed using the IncuCyte software. In parallel (right graph), the same cell-cell fusion was performed in impedance E-plates and the CEI was recorded in real-time, starting from the time point of cell overlay. The graph depicts the impedance signal (expressed as cell index) over time of 4 technical replicates (mean ± SD).

